# Tecovirimat is highly efficient on the Monkeypox virus lineage responsible for the international 2022 outbreak

**DOI:** 10.1101/2022.07.19.500484

**Authors:** Gaëlle Frenois-Veyrat, Franck Gallardo, Olivier Gorgé, Elie Marcheteau, Olivier Ferraris, Artem Baidaliuk, Anne-Laure Favier, Cécile Enfroy, Xavier Holy, Jérémy Lourenco, Rhéa Khoury, Flora Nolent, Douglas W. Grosenbach, Dennis Hruby, Audrey Ferrier, Frédéric Iseni, Etienne Simon-Loriere, Jean-Nicolas Tournier

## Abstract

The ongoing monkeypox virus (MPXV) outbreak is the largest ever recorded outside of Africa. Genomic analysis revealed a divergent phylogenetic lineage within clade 3, and atypical clinical presentations have been noted. We report the sequencing and isolation of the virus from the first clinical case diagnosed in France in May 2022. We tested the *in vitro* effect of tecovirimat (ST-246), a FDA approved drug, against this novel strain, showing efficacy at the nanomolar range. In comparison, cidofovir showed activity at micromolar concentrations. These results and the safety profile of tecovirimat strongly support its use in clinical care of severe forms for the 2022 MPXV outbreak.

## Main text

Monkeypox virus (MPXV) is a zoonotic virus belonging to the Orthopoxvirus genus, which also includes vaccinia virus and variola virus^1^. Since May 2022, MPXV is causing the largest human outbreak ever recorded outside of Africa^2^. Recent studies report atypical demographics and clinical findings in cases of MPXV, mainly but not exclusively found in men who have sex with men^3, 4, 5^. The first suspected case in France was a male patient consulting at an HIV and sexual health center on May 19^th^, 2022 for vesiculopustular lesions on the face and the genital organs, with no reported travel history. At the clinical examination, the patient presented a lesion on the lip, the nipple and two lesions on the genitalia. The MPXV infection at all three sites was confirmed by qPCR assays at the National Reference Center for orthopoxviruses. The PCRs, adapted from previous studies, consisted in a first screening for the genus orthopoxvirus^6^, followed by a second, specific, qPCR for MPXV diagnosis^7^. The viral strain MPXV/France/IRBA2211/2022 was isolated on Vero cells (see methods) and is available upon request, as it may be useful to evaluate diagnostic tools, antivirals or vaccines. Transmission electron microscopy was performed on infected Vero cells. The resulting image (Figure 1A) showed classical pox-like intracellular virions of approximately 250 nm × 125 nm in size with an hourglass-shaped core. The genome from both the primary samples and the isolate were identical, and clustered within the newly proposed B.1 lineage within clade 3^8^ (formerly designated as the “West African” clade^9^) (Figure 1B).

**Figure 1.**
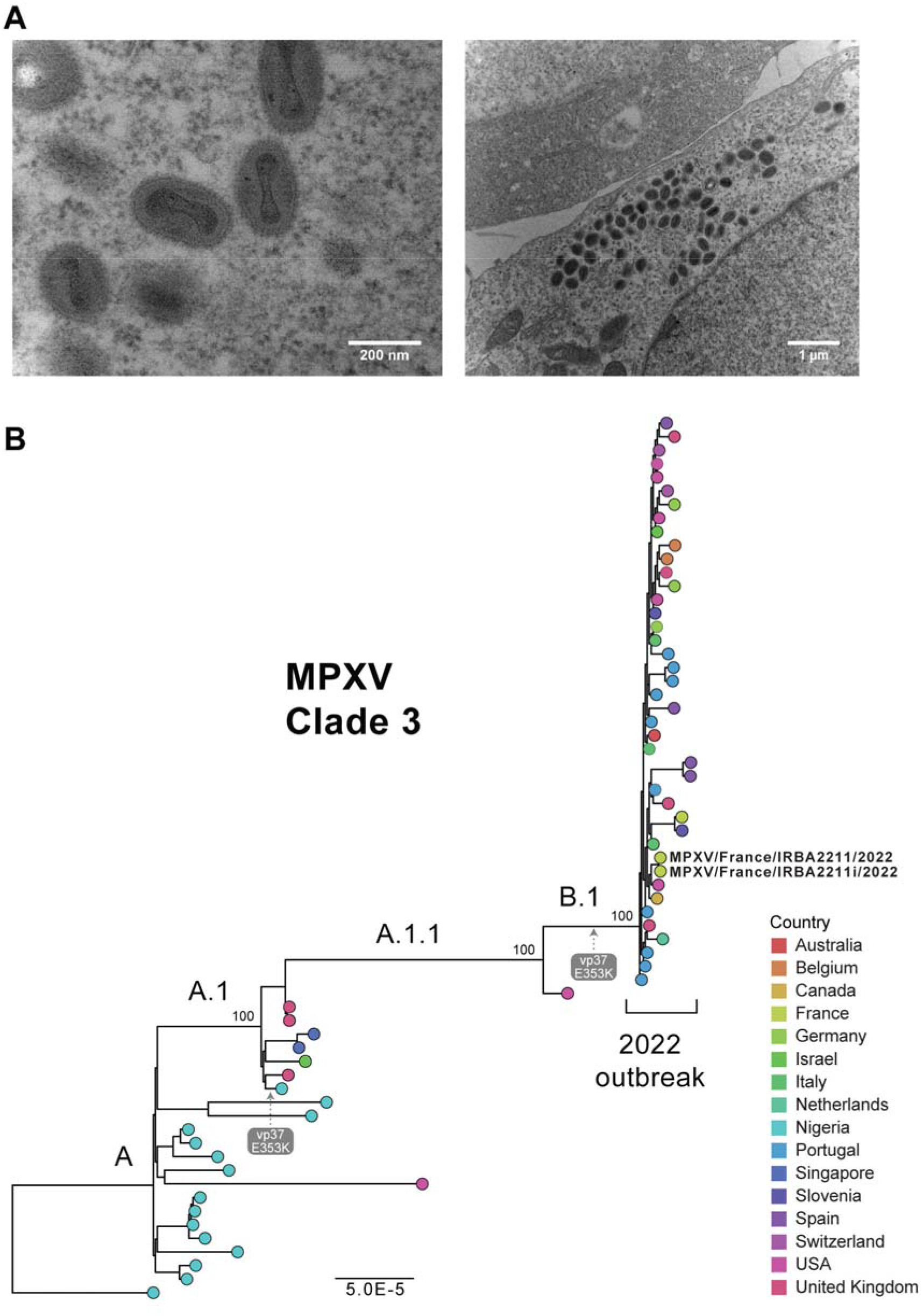
A. Transmission electron microscope view of MPXV virions (MPXV/France/IRBA2211/2022) within cells captured at 48h post infection, at higher (left) and lower (right) magnification. B. Maximum likelihood phylogenetic tree of MPXV Clade 3 contains sequences from previous endemic circulation in Nigeria, and later outbreaks in other countries, including the global outbreak in 2022 (subclade B.1). The tree is rooted according to the full phylogeny of MPXV and clade support values are shown as percentages above the corresponding branches. The tree tips are colored by country and sequences from this study marked with strain names. E353K label marks a mutation in the F13L homolog, occurring in all sequences of B.1 subclade and 1 subclade A sequence (MT903341).

MPXV infection leads to a wide spectrum of clinical presentation and disease severity, including pauci- or asymptomatic infection as reported within the 2022 outbreak^10^. The majority of the clinical characteristics of human MPXV infection mirror those of smallpox, although with a milder mortality rate varying from 10 % for clade 1 (former Congo Basin clade), to 3.6 % for clades 2 and 3. Human MPXV infection often begins with a combination of non-specific symptoms such as fever, chills, asthenia, lymph node swelling, back pain, and muscle aches. This prodromal phase is followed by the classical rash phase evolving swiftly in papules turning into vesicles and pustules, evolving into scabs. The number of skin lesions may range from a few to thousands. The current outbreak of MPXV infection is characterized by several atypical features such as shorter incubation phase^11^, and milder clinical presentation often associated with localized rash and lesions in genital and perianal sites, sometimes highly painful^3, 4, 5^. Importantly, severe disease can occur in immunocompromised patients, pregnant women, and children.

No specific treatment is currently approved for MPXV infection. However, several molecules developed to treat smallpox (Tecovirimat (ST-246); Cidofovir; Brincidofovir) have shown broad spectrum activity both *in vitro* and in animal models against multiple orthopoxviruses including MPXV, variola virus and rabbitpox^12, 13, 14^. While brincidofovir treatment of 3 MPXV infected patients between 2018 and 2021 was rapidly stopped due to elevated liver enzymes^15^, no safety signals were identified in reports of compassionate use of tecovirimat, making it a potential resource to help patients in the current outbreak. Tecovirimat has been authorized in Europe by the European medicine agency under ‘exceptional circumstances’, although its efficacy has not been shown on the current circulating strain^16^. Tecovirimat has been shown to target vaccinia virus (VACV) F13L gene product VP37 protein, required for extracellular virus particle formation. The protein is highly conserved within the orthopoxvirus genus. Interestingly, all 2022 MPXV genomes carry the E353K substitution in the VP37 protein, which was absent from the most recent common ancestor within clade 3 (Figure 1B), but noted in a single 2018 outbreak genome (MT903341).

To assess the activity of Tecovirimat, we performed doses response studies on Vero cells (see methods). Tecovirimat completely abolished MPXV production above 100 nM, with an IC_50_ of 12.7 nM. For comparison, we used a VACV ANCHOR™-GFP infection model, where virus infection and propagation can be visualized in living cells, revealing a comparable IC_50_(6 to 8.6 nM) (Figure S1). Cidofovir IC_50_ on MPXV lineage B.1 was 30 μM (Fig. 2), 3,000 less potent than tecovirimat *in vitro.*

**Figure 2:**
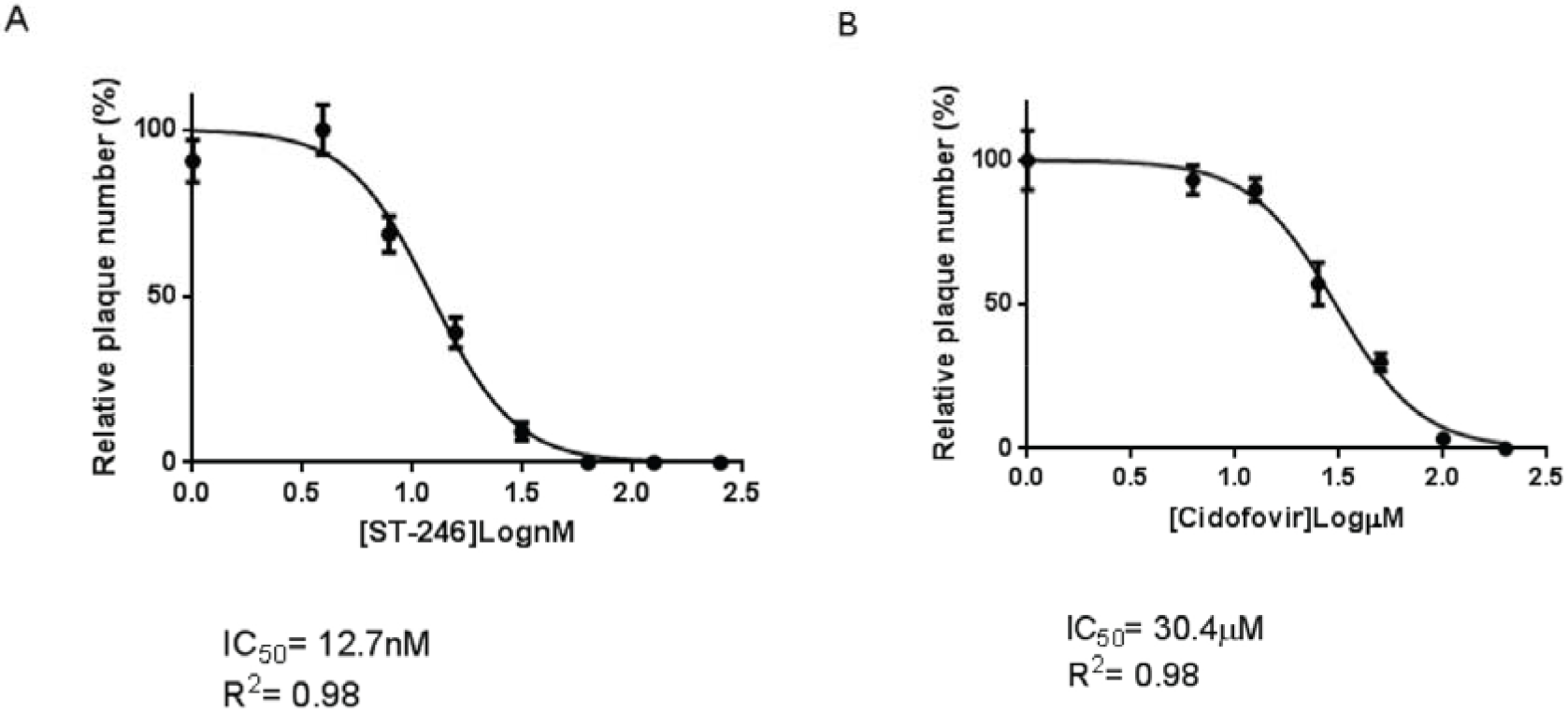
ST-246 and Cidofovir inhibits plaque formation of the MPXV MPXV/France/IRBA2211/2022 isolate. A) Vero cells are infected with MPXV/France/IRBA2211/2022 and treated with indicated concentrations of ST-246 (A) or Cidofovir (B) for 72h. Lysis plaque inhibition is expressed in % normalized over CTL conditions. IC_50_ and R^2^ are indicated.

These results support the use of tecovirimat in the clinical response to the 2022 MPXV outbreak, in particular for immunosuppressed patients. Of note, tecovirimat is not recommended for pregnant women due the absence of safety data.

Considering that resistance mutations to tecovirimat under artificial culture pressure have been described for various orthopoxviruses^17^ (Figure S2), attention to adherence to treatment will be needed, especially in the context of milder clinical forms, and it will be important to monitor of the evolution of the virus during its outbreak circulation.

## Methods

### Ethics

This work has received patient consent through the national reference center for orthopoxviruses (CNR-LE orthopoxvirus).

All work with infectious virus was performed in BSL-3 containment laboratories.

### Virus detection

Test and control samples were extracted using the QIAamp DNA Mini kit (Qiagen) with small modifications. Briefly, pre-extraction inactivation was performed in BSL3 by adding 200μLof sample to a tube containing 200 μL of Qiagen Buffer AL and 20 μL of proteinase K. The tubes were incubated at 70°C for 10 min to inactivate the virus prior to extraction.

The *pan-Orthopoxvirus* and pan-MPXV PCR assay have been previously described^6, 7^. Each reaction consisted of 5 μl of extracted DNA and 15 μL of iTaq™ Universal Probes Supermix (BioRad), containing 0.4 mM of each primer, 0.25mM of probe in the *pan-Orthopoxvirus* PCR, and 0.2mM of probe in the pan-MPXV PCR. All assays were performed on a CFX96 thermocycler (BioRad). Data and results were analyzed and reported using Bio-Rad CFX Maestro.

### Virus isolation, titration and production

The swabs used for sample collection were discharged in universal transport media. After homogenization, the samples were diluted in Dulbecco’s Modified Eagle Medium (DMEM) containing 0.4 % fetal bovine serum (FBS). To inhibit bacterial and fungal contamination antibiotics and antifungal were added (Gentamycin: 2,5μg/ml, Mycostatin: 10 U/ml, Penicillin: 100 U/ml, Streptomycin: 100 μg/ml). Following clarification at 700 g for 10 min, we inoculated 50 μL of the sample onto monolayers of Vero (African green monkey kidney, ATCC CCL-81) cells in 96 well plates for both titration and isolation (eight replicates / dilution). The plates were incubated at 37°C under 5 % CO_2_ and observed daily for potential cytopathic effect. The viral titer was calculated according to the Reed and Muench method and expressed in terms of TCID_50_/mL^18^.

Supernatants from wells presenting cytopathic effect were harvested and inoculated on T25 flasks. At day 3, upon observation of 80 % cytopathic effect, the flasks were frozen/thawed 3 times and the production was clarified at 1,200 g for 10 min (passage 2). Passage 2 stock was tittered by plaque forming assay in 24-well plates and sequenced. The titer was 4.1×10^6^ PFU/mL

### Transmission electron microscopy

Two days after infection, cells were fixed with 2.5 % (vol/vol) glutaraldehyde in sodium cacodylate buffer (0.1 M; pH 7.4; 10 mM CaCl2, 10 mM MgCl2, and 2% glucose) for 4 days at 4°C. After washing samples with a mixture of saccharose (0.2 M) and sodium cacodylate (0.1 M), cells were postfixed them with 1% (vol/vol) osmium tetraoxide in cacodylate buffer for 1 h at room temperature. Samples were stained with 2% (vol/vol) uranyl acetate for 1 h at 4°C followed by gradual dehydration with increasing ethanol concentrations. Samples were embedded in Epon LX112 resin (Ladd Research Industries) in embedding capsules and polymerized for 24 h at 60°C. Embedded samples were cut in ultrathin 80-nm sections with an UC6 ultramicrotome (Leica), placed them onto 300-mesh copper grids, stained sections with 2 % uranyl acetate and lead citrate, and examined them under a Philips CM10 TEM microscope (operating at 100 kV) and equipped with a Denka LaB6 cathode and a CCD Erlanghsenn 1000 Gatan camera. No filtering procedures were applied to the images.

### Sequencing

Viral DNA was extracted from the clinical isolate or cell supernatant using the Macherey Nagel pathogen kit according to the manufacturer’s instructions after DNase treatment. Libraries were prepared in parallel for Oxford Nanopore Technologies (ONT) and Illumina. For ONT, we used the ligation sequencing kit (SQK-LSK109) after whole genome amplification (Cytiva kit, FisherScientific), following manufacturer’s instructions.

Data were filtered with kraken/bracken (v2.1.2/v2.6.2) to identify reads matching the chordopoxvirinae family. We mapped reads to Monkeypox virus strain Israel_2018 (MN648051.1) with minimap2 v2.24. From this reads mapping, a consensus sequence was extracted using nanopolish (v 0.13.2) and bcftools (v1.14). We obtained an average coverage depth of 620X (53,632 mapped reads out of 130,176).

In parallel, libraries were prepared for Illumina sequencing using the NEBNext Ultrall DNA kit (New England Biolabs), after shearing with an M220 Covaris device (Covaris) following the manufacturer’s recommendations. Non-amplified (“Native”) as well as amplified extracts were processed. After size selection with AMPure beads (Beckman), sequencing was performed on a NextSeq 550 instrument (Illumina) using a HT 2×150 bp cartridge.

We mapped the reads with bwa mem (v0.7.17-r1188) and obtained two datasets with a marked difference between native and amplified samples. We obtained 46.2 % of mapped reads for MPXV/France/IRBA2211/2022 after amplification, and only 9.5 % for the native extract. We attributed the difference to the additional DNase treatment performed for amplified samples. There was no nucleotide difference among the two sequences, indicating that amplification did not alter original sequences.

We then aggregated the sequencing data from the two platforms by using the Nanopore consensus obtained from minimap2 as reference for mapping the Illumina reads. We filtered the sam file to remove unmapped reads and used Geneious R11 (Biomatters) to extract a consensus sequence (ON755039).

### Phylogenetic analysis

All available MPXV sequences were retrieved from GenBank on July 3^rd^, 2022. This dataset was subset to keep published and discussed sequences^19^ belonging to the newly proposed clade 3. We used Nextalign as implemented in the MPXV Nextstrain^20^ build to align sequences, also adopting their masking strategy (https://github.com/nextstrain/monkeypox). The alignment was visually inspected for accuracy. A maximum likelihood phylogeny was inferred with IQ-TREE v.2.0.6^21^ with 1,000 (ultrafast) bootstrap^22^ replicates for branch support estimates. The tree was rooted between MPXV strain Nigeria-SE-1971 (KJ642617) and sequences of subclade A in accordance with the full comprehensive MPXV phylogenetic tree.

### F13L homolog alignment

F13L homologs sequences were extracted from the available full genomes orthopoxviruses tested for susceptibility to tecovirimat: VACV (AY243312.1), VARV (DQ441447.1), CPXV (AF482758.2), CMLV (AY009089.1), MPXV (HM172544.1), ECTV (AF012825.2)^23^, as well as RPXV (AY484669.1)^24^. Amino acid sequences were aligned with MAFFT v7.450^25^. Annotations for mutations associated with resistance to tecovirimat^26^ were added to the alignment plot in Geneious Prime (Biomatters).

### Antiviral assays

For MPXV infections, Vero cells were seeded in six-well plates (1.5 10^6^ cells/well). Twenty-four hours post seeding, cells were infected with MPXV/France/IRBA2211/2022 in order to obtain 50-100 pfu/well. After adsorption for one hour, the inoculum was replaced with DMEM and 2 % FCS containing the appropriate concentration of the drug ST-246. After one hour at 37°C, cells were then overlaid with 1.6 % carboxymethyl cellulose (VWR) diluted in DMEM and 2 % FCS. The plates were incubated for three days at 37°C in a 5 % CO2 incubator. Monolayers were fixed and stained in 3.7 % formaldehyde, 0.1 % crystal violet and 1.5 % methanol. The plaques were counted microscopically. All experiments were performed in triplicate.

For VACV ANCHOR™ infections, HeLa cells (ATCC CCL-2) were grown in DMEM without phenol red (Sigma Aldrich) supplemented with 10 % FBS (Eurobio-Scientific), 1mM sodium pyruvate (S8636; Sigma Aldrich), L-Glutamine (G7513; Sigma Aldrich) and Penicillin-streptomycin solution (P0781; Sigma Aldrich). Cells were plated (9,000 cells per well) in Corning Cellbind 96 well plates in complete DMEM. Twenty-four hours post-seeding, cells were treated with the test compound at the indicated concentration, in triplicate. Cells were then infected with VACV ANCHOR™ (Copenhagen strain with the ANCHOR system in the TK locus) virus at MOI 0,1 and incubated at 37°C and 5 % CO_2_. At 48 h and 72 h, cells were fixed with 4 % formalin (Sigma) for 10 min at RT, washed with PBS and incubated with PBS-Hoechst 33342 (1 mg/mL).

Data acquisition by high content microscopy was performed on a Thermo Celllnsight CX7 HCS microscope using a compartmental analysis algorithm. Results were extracted, normalized over the vehicle-treated condition, and expressed as the average of three independent wells +/-SD. CC50 were calculated using GraphPad Prism v9.

## Supplementary Figure legends

**Figure S1:**
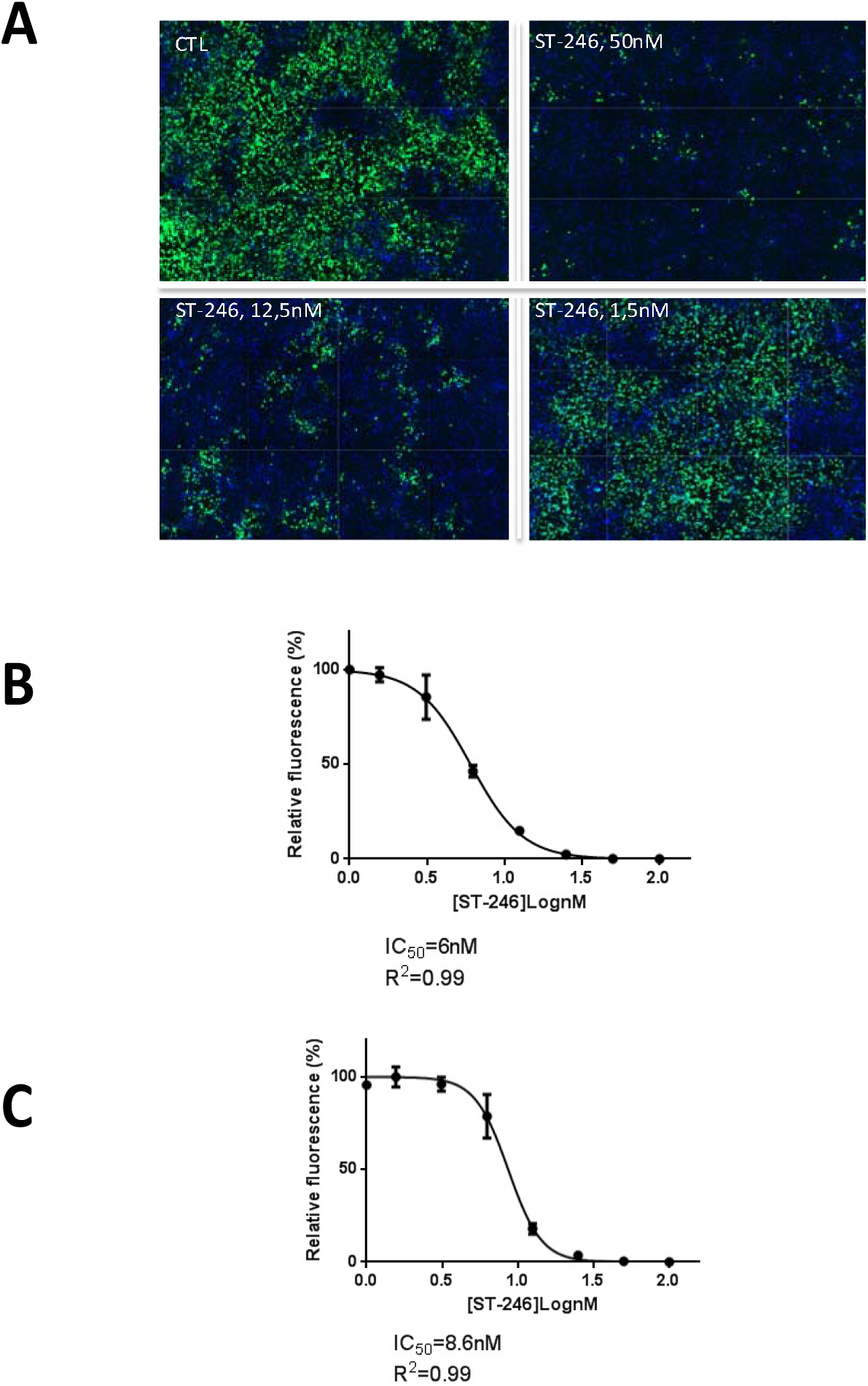
ST-246 inhibits VACV ANCHOR™-GFP propagation at nM concentration. A) Reconstruction of 12 imaging fields captured on a CX7 Celllnsight HCS microscope 48h post infection with indicated concentration of ST-246. Blue: nuclei, green VACV ANCHOR™ B/C) Quantification of the impact of ST-246 on the number of infected cells at 48 h (B) and 72 h(C) post-infection. IC_50_ and R^2^are indicated.

**Figure S2:**
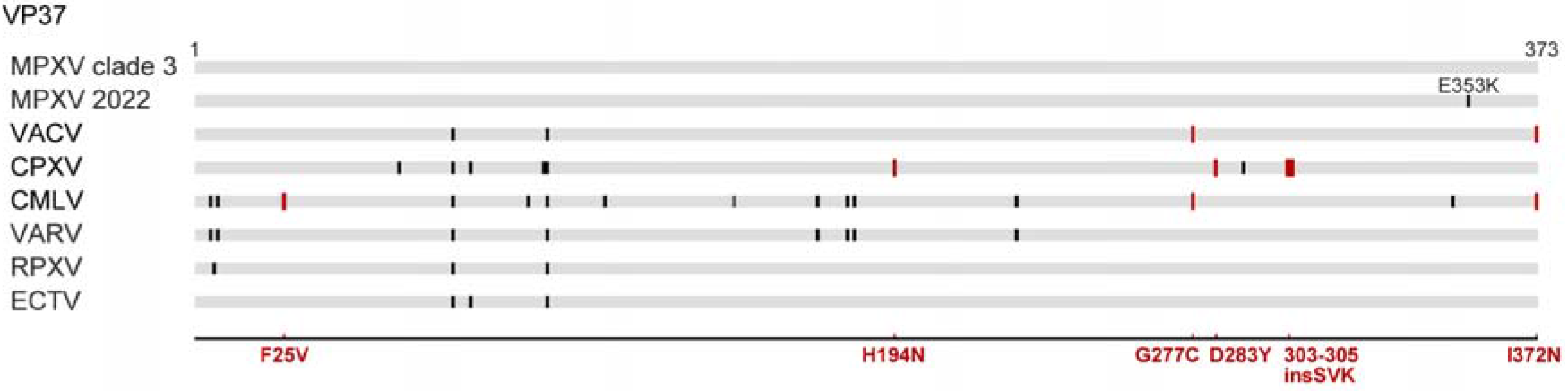
VP37 (F13L homologs) amino acid alignment to MPVX clade 3 shows multiple differences, marked with black vertical bars, with other orthopoxviruses susceptible to tecovirimat. The E353K mutation in VP37 is a signature substitution of MPXV lineage B.1 (2022). Experimentally generated and verified mutations associated with resistance to ST-246 are marked in red below the alignment and well as in the corresponding OPV species in which they have been experimentally tested (vertical red bars). Importantly, these mutations (in red) are annotated in the alignment in the corresponding positions, while they do not exist in the original sequences.

**Figure S3:**
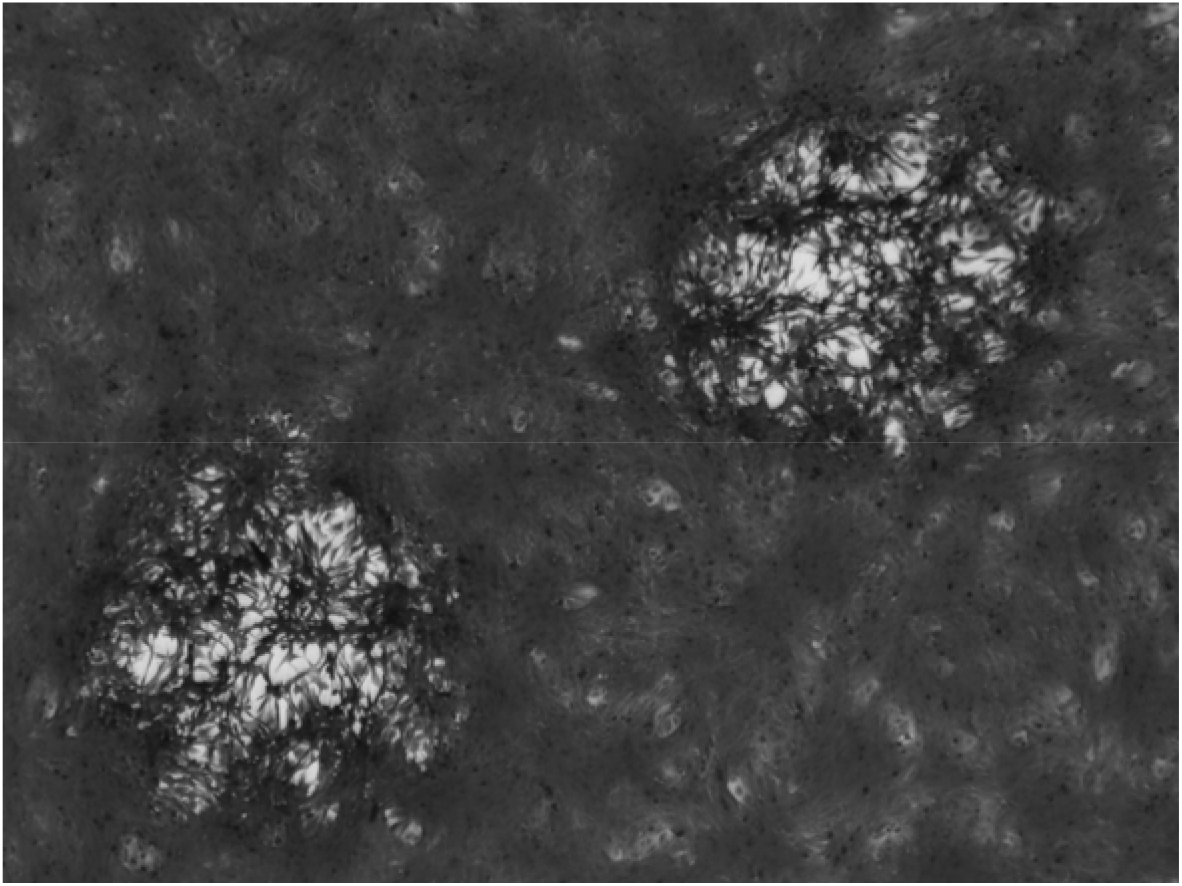
Cytopathic effect on Vero cell line of MPXV/France/IRBA2211/2022 strain (optical microscopy 20x).

## Data availability

All data supporting the findings of this study are available in the manuscript. The MPXV genomic data analysed here are available on GenBank (https://www.ncbi.nlm.nih.gov/genbank/, ON755039).

## Acknowledgments

We would like to thank all the healthcare workers, public health employees, and scientists involved in the MPX outbreak response. We would like to acknowledge the technical skills of Cedric Casteliarin for preparing the samples of transmission electron microscopy. We acknowledge the authors, originating and submitting laboratories of the orthopoxviruses sequences deposited on GenBank. This work used the computational and storage services provided by the IT department at Institut Pasteur, Paris.

## Author contribution

Conception and design of experiments: F.I., J-N.T., E.S-L.; Patient care: J.L., R.K.; Virus isolation, production, titration: O.F., A.F-R., G.F-V; Electron Microscopy: A-L.F., C.E., X.H.; Sequencing: O.G., F.N.; MPXV antiviral assay: F.I., G.F-V.; VACV antiviral assay: F.G., E.M.; Antiviral material: D.W.G., D.H.; Phylogenetic analysis: A.B. and E.S-L.; Writing – original draft, review and editing: J-N.T. and E.S-L. with inputs from all authors.

